# IRE1-mediated degradation of *pre-miR-301a* promotes apoptosis through upregulation of *GADD45A*

**DOI:** 10.1101/2023.06.21.545854

**Authors:** Magdalena Gebert, Sylwia Bartoszewska, Lukasz Opalinski, James F. Collawn, Rafał Bartoszewski

**Author notes:** Correspondence: Rafał Bartoszewski, Department of Biophysics, Faculty of Biotechnology, University of Wrocław, F. Joliot-Curie 14a Street, 50-383 Wrocław, Poland.

## Abstract

The unfolded protein response is a survival signaling pathway that is induced during various types of ER stress. Here we focus on the IRE1 pathway to determine IRE1’s role in miRNA regulation during ER stress. During induction of ER stress in human bronchial epithelial cells, we utilized next generation sequencing to demonstrate that *pre-miR-301a* and *pre-miR-106b*, were significantly increased in the presence of an IRE1 inhibitor. Conversely, using nuclear-cytosolic fractionation on ER stressed cells, we found that these three pre-miRNAs were decreased in the nuclear fractions without the IRE1 inhibitor. We also found that *miR-301a-3p* targets the proapoptotic UPR factor, growth arrest and DNA-damage-inducible alpha (*GADD45A*). Inhibiting *miR-301a-3p* levels or blocking its predicted miRNA binding site in *GADD45A*’s 3’ UTR with a target protector increased *GADD45A* mRNA expression. An elevation of *XBP1s* expression had no effect on *GADD45A* mRNA expression. We also demonstrated that the introduction of a target protector for the *miR-301a-3p* binding site in *GADD45A* mRNA during ER stress promoted cell death in the airway epithelial cells. These results indicated that IRE1’s endonuclease activity is a two-edged sword that splices *XBP1* mRNA for survival and degrades *pre-miR-301a* to elevate the mRNA expression of a pro-apoptotic gene*, GADD45A*.

## Introduction

Endoplasmic reticulum (ER) homeostasis is dependent upon proper folding and maturation of secretory pathway proteins, lipid biosynthesis, and cellular calcium homeostasis. Disruption of the ER homeostasis, however, leads to a stress response called the unfolded protein response (UPR) that is a multifunctional signaling pathway (1–3). The UPR serves primarily as a cellular adaptive mechanism that counteracts the stress-related deregulation of ER function and promotes cellular survival from both intrinsic and extrinsic insults (4,5). If the recovery mechanisms are ineffective, however, and the ER stress is maintained, the UPR can promote cell death (6–10).

The UPR is mediated by three proximal ER stress sensors that include the inositol-requiring enzyme 1 alpha (IRE1), the PKR-like endoplasmic reticulum kinase (PERK), and the activating transcription factor 6 (ATF6) (6–10). Among them, IRE1 regulates the phylogenetically most conserved arm of the UPR that is involved in the balance between cell survival and cell death (11–13). IRE1’s unique feature is its UPR-related RNase activity that is responsible for the production of a potent transcription factor, X box-binding protein-1 (XBP1s). IRE1 regulates this activation through the nonconventional splicing of *XBP1* mRNA and the degradation of a large pool of cytoplasmic mRNAs in a process called regulated-IRE1 dependent decay (RIDD) (9,14). The complex role of IRE1 in the regulation of mammalian UPR cannot be fully explained by IRE1’s one known, specific RNA target, *X box-binding protein-1* (*XBP1*) or through the known mRNA substrates of its RIDD activity. Here, we focused on the role that IRE1 plays on miRNA expression during the UPR.

Using next-generation sequencing (NGS) of normal immortalized human bronchial epithelial cells, 16HBE14o-, we identified three miRNAs, *hsa-miR-301a-3p*, *hsa-miR-106b-5p*, and hsa*-miR-17-5p* that had decreased levels of expression during the UPR. Our analyses demonstrated that all 3 of their pre-miRNA precursors were degraded by IRE1. Furthermore, we demonstrated that the IRE1-mediated downregulation of *hsa-miR-301a-3p* allowed for the accumulation of a proapoptotic UPR factor, growth arrest and DNA-damage-inducible alpha (*GADD45A*), and this is known to contribute to cell fate decisions (15–17). The studies presented here demonstrate that during UPR, IRE1 degrades *pre-miR-301a* and this enhances the expression of *GADD45A*, and subsequently drives the ER stressed cells towards apoptosis. This study demonstrates how inhibition of miRNA maturation by IRE1 can modulate cell fate decisions during the UPR.

## Materials and Methods

### Cell culture

Human bronchial epithelial 16HBE14o- cells (Sigma-Aldrich, Cat. #SCC150, Poznan, Poland) were obtained as previously described (15,18,19). 16HBE14o- cells were cultured in minimum essential modified Eagle’s medium (Invitrogen, Carlsbad, CA, USA) with 10% FBS in a humidified incubator at 37°C in 5% CO_2_ in 6-well plates and allowed to grow to 70–80% confluence prior to the start of the experiments. The inducible stable HeLa S3 *XBP1s* cell line containing the cDNA sequence of *XBP1s* (NM_001079539.1) was obtained as described previously (20). This cell line was cultured in Minimum Essential Modified Eagle’s Medium with 2 mM glutamine, hygromycin B (300 µg/ml) and 10% tetracycline free fetal bovine serum (Takara Bio, USA) in a humidified incubator at 37°C in 5% CO_2_ in 6-well plates. For the XBP1s 24 h induction, doxycycline (400 µg/ml, D3072 Millipore-Sigma) was added to culture media for 24 hours. Cells were allowed to grow to 70–80% confluence before the start of the experiments. The Ambion™ PARIS™ system was used for the isolation of both RNA and native protein from the same experimental sample (AM1921) and was used according to the manufacture’s protocol to obtain the nuclear and cytosolic fractions. The follow up isolation of RNA and small non-coding RNA was performed with miRNeasy kit (Qiagen).

#### Induction of ER stress and activation of the UPR

Pharmacological induction of ER stress and activation of the UPR were performed as previously described (15,19,21,22). Briefly, cells were treated with the compounds for the time periods specified: Tm (2.5 μg·mL^−1^; Sigma, T7765), Tg (50 nm; Sigma, T9033), calpain inhibitor I (ALLN; 100 μm; Abcam, ab146608, Cambridge, UK). CTRL cells were treated with vehicle CTRL, DMSO (< 0.5% v/v; Sigma, D2650). To verify IRE1 activity, cells were treated with 20 µM 4µ8C (an IRE1 inhibitor, Sigma-Aldrich, SML0949) dissolved in DMSO (Sigma-Aldrich, St. Louis, MI, USA) (20,21,23).

#### Isolation of RNA and small non coding RNA

Total RNA (containing both mRNA and miRNA) was isolated using miRNeasy kit (Qiagen). RNA concentrations were calculated based on the absorbance at 260 nm. RNA samples were stored at −70°C until use.

#### miRNA analogs and target protector transfections

Cells were seeded onto 6-well plates or 35-mm dishes and transfected at 70–80% confluence with Lipofectamine RNAiMax (Thermo Fisher Scientific) according to the manufacturer’s protocol. mirVana miRNA Mimics and mirVana miRNA Inhibitors (both from Thermo Fisher Scientific) were used at final concentrations of 30 and 150 nM, respectively. mirVana pre-miRs and inhibitors used in this study are as follows: *pre-miR-106b-5p* (MH10067), *pre-miR-301a-3p* (MH10978) and inhibitors: *miR-106b-5p* (PM10067) and *miR-301a-3p* (MC10978) and inhibitor (PM10978). As additional controls, *mir*Vana™ miRNA Mimic Negative Control #1 (4464058), and *mir*Vana™ miRNA Inhibitor Negative Control #1 (4464076) were used. The Ambion Silencer Negative Control No. 1 siRNA (4390843; Thermo Fisher Scientific) was used as well.

The target protector (TP) and the respective control (morpholino oligo that targets a human beta-globin intron mutation that causes beta-thalassemia, 5’ -*CCTCTTACCTCAGTTACAATTTATA*-3’) were purchased from Gene Tools. LLC (Philomath, OR, USA) and directed against the *hsa-miR-301a-3p* binding site in *GADD45A (*5′-*ACTTCAGTGCAATTTGGTTCAGTTA-3’*) mRNA. TPs were used at a final concentration of 1 µM. The transfected cells were cultured for 2 days before analysis.

### Measurement of mRNA quantitative Real Time PCR (qRT-PCR)

We used TaqMan One-Step RT-PCR Master Mix Reagents (Applied Biosystems) as described previously (19,24) using the manufacturer’s protocol (retrotranscription: 15 min, 48°C). The relative expressions were calculated using the 2^-ΔΔCt^ method (25) with the *RPLP0 (Hs00420898_gh)* gene as reference genes for the mRNA and *RNU44* (001094) or *RNU6 (*HS_RNU6-2_11) for the miRNA and pre-miRNA, respectively. TaqMan probes ids used were: *BIP* (Hs00607129_gH), *GADD45A* (Hs00169255_m1), *MT-CYB* (Hs02596867_s1), *hsa-miR-106b* (000442), *hsa-miR-301a* (000528), *hsa-miR-17* (002308), *MT-CYB* (Hs02596867_s1), *pre-miRNA-17* (HS_miR-17_1_PR, MP00001064), *pre-miRNA-106b* (HS_miR-106b_1_PR, MP00000147), *pre-miRNA-301a* (HS_miR-301a_1_PR, MP00001771), *pri-miR-17* (Hs03295901_pri), *XBP1(S) (*Hs03929085_g1). *EREG* (Hs00914313_m1).

### Measurement of RNA half-life. *GADD45A*

*hsa-miR-301a-3p*, *hsa-miR-106b-5p*, *pre-miR-301a* and *pre-miR-106b* RNA half-lives were measured as described in (26,27) with the following modifications. Cells were grown on 35-mm plastic dishes to ∼80% confluency. Parallel experiments were performed in cells exposed to 2.5 µg/ml Tm, Tm and 20 µm 4µ8C, or cultured in control conditions. Actinomycin D (5 μg/ml; MilliporeSigma, Burlington, MA, USA) was added to stop transcription at the start of the experiment, and the RNA was isolated at the indicated time intervals using Qiagen miRNeasy. Actinomycin D was maintained throughout the experiments. Total RNA levels were measured by real-time PCR at each time point using TaqMan-based assays and normalized to endogenous 18S rRNA levels (amplified using a standard primer set). RNA values for each time point were calculated from 3 individual samples generated in at least 2 independent experiments. Relative RNA levels were plotted as differences from RNA levels at the initial time point (t = 0). The RNA half-lives were calculated from the exponential decay using the trend line equation C/C_0_ = e^−kdt^, where C and C_0_ are RNA amounts at time t and t_0_, respectively, and k_d_ is the RNA decay constant as previously described (28,29).

### Western Blots

Cells were lysed in SDS lysis buffer (4% SDS, 20% glycerol, 125 mM Tris-HCl pH=6.8) supplemented with protease inhibitors (cOmplete ™ Mini (Roche)). The insoluble material was removed by centrifugation at 15,000 g for 15 min. Protein concentrations were determined by Bio-Rad™ DC-Protein Assays using bovine serum albumin (BSA) as standard. Following the normalization of protein concentrations, the lysates were mixed with an equal volume of 2X Laemmli sample buffer and incubated for 5 min at 95°C prior to separation by SDS PAGE on Criterion TGX stain-free 4-15% gradient gels (Bio-Rad). Following SDS-PAGE, the proteins were transferred to polyvinylidene fluoride membranes (300 mA 4 hours at 4°C). The membranes were then blocked with BSA (Sigma-Aldrich) dissolved in PBS/Tween-20 (3% BSA, 0.5% Tween-20 for 1-2 hours), followed by immunoblotting with the primary antibody: rabbit anti-GADD45A (1:2000, D17E8; Cell signaling); rabbit anti-XBP-1 (S) (1:1000, D2C1F, Cell Signaling). After the washing steps, the membranes were incubated with goat anti-rabbit IgG (H+L chains) HRP-conjugated secondary antibodies (Bio-Rad) and detected using SuperSignal West Pico ECL (Thermo Fisher Scientific). Densitometry was performed using Image Lab software v. 4.1 (Bio-Rad).

#### Real-time cell viability assay

For real-time monitoring of cell viability, we applied real time and label free holographic microscopy-based monitoring of cell death and viability using HoloMonitor M4**®** time-lapse cytometer (Phase Holographic Imaging PHI AB, Lund Sweden). Holographic microscopy was used to follow the optical thickness and irregularity of cells exposed for up to 24 hours to Tm in the presence or absence of *pre-miR-301a*. The images from up to 5 independent optical fields were collected and analyzed according to manufacture instructions with HoloMonitor® App Suite software. Healthy cells are irregular in shape and thin, and dying cells are round and thick (30–35). For all analysis, the same cells parameters qualification was applied.

#### Next-generation RNA sequencing analyses

The RNA isolation and analyses were performed in 16HBE14o- cells as previously described (19). Briefly, following total RNA isolation, samples were validated with qRT-PCR for ER stress activation prior to further analysis. Following rRNA depletion, the remaining RNA fraction was used for library construction and subjected to 100-bp paired-end sequencing on an Illumina HiSeq 2000 instrument (San Diego, CA, USA). Sequencing reads were aligned to the human reference genome assembly (hg19) using TopHat (36). Transcript assembly and estimation of the relative abundance and tests for differential expression were carried out with Cufflinks and Cuffdiff (37). The resulting data were validated with qRT-PCR. The heat map generation and hierarchical clustering were performed with the Morpheus Web server (Morpheus, https://software.broadinstitute.org/morpheus).

Small RNA sequencing libraries were prepared using QIAseq miRNA library kit (Qiagen) following the manufacturer’s instructions and followed by sequencing on an Illumina NextSeq instrument. Using Qiagen’s Gene Globe Software, sequencing reads were aligned to the human reference genome assembly (hg19) followed by transcript assembly and estimation of the relative abundances. The analysis of the differential expression of small RNAs between control and experimental samples were performed with geNorm (38) in the Gene Globe Software.

### Statistical analysis

Results were expressed as a mean ± standard deviation. Statistical significance was determined using the Student’s t-test with P<0.05 considered significant. The correlation was accessed via the Pearson product-moment correlation method.

## Results

To gain insight into IRE1’s role as a direct miRNA regulator during ER stress, we followed next generation sequencing changes in miRNA levels in 16HBE14o- cells that were exposed for 6 hours to ER stress (induced by ALLN or tunicamycin (Tm)) in the presence of an IRE1 specific inhibitor, 4µ8C (39). In previous studies, we used a 6-hour time point as putative transitional time point for the cell fate decision process (15,19). We focused this current analysis on miRNAs that were downregulated in both ER stress models, and that their sequence contained the consensus IRE1 target sequence (UGCA) (40). These two criteria would make them potentially prone to IRE1 endoribonuclease activity. In the case of ALLN treated cells, all of the selected miRNAs were induced when IRE1 activity was impaired. In Tm-treated cells, however, their miRNA levels were more resistant to UPR-related reduction, and were only slightly reduced or unaffected (**Figure 1A**). Furthermore, as shown in **Supplemental Figure 1**, IRE1 inhibition in tunicamycin-treated cells resulted in elevated levels of 12 miRNAs, whereas the levels of 51 were reduced. Interestingly, the presence of 4µ8C during ALLN treatment increased the expression of 162 miRNAs and reduced levels of 964 others when compared to the “no stress” exposed cells in the presence of IRE1 inhibitor. The observed discrepancies in the miRNA profiles between these two models resulted presumably from the different mechanisms of action of the two stressors, including inhibitory effects of ALLN on the IRE1 branch (15,21). Consequently, we limited the use of this NGS data to support initial selection of IRE1-dependent miRNAs and verified a potential positive control that was previously reported as an IRE1-degraded miRNA, *miR-17-5p* (41).

**Figure 1.**
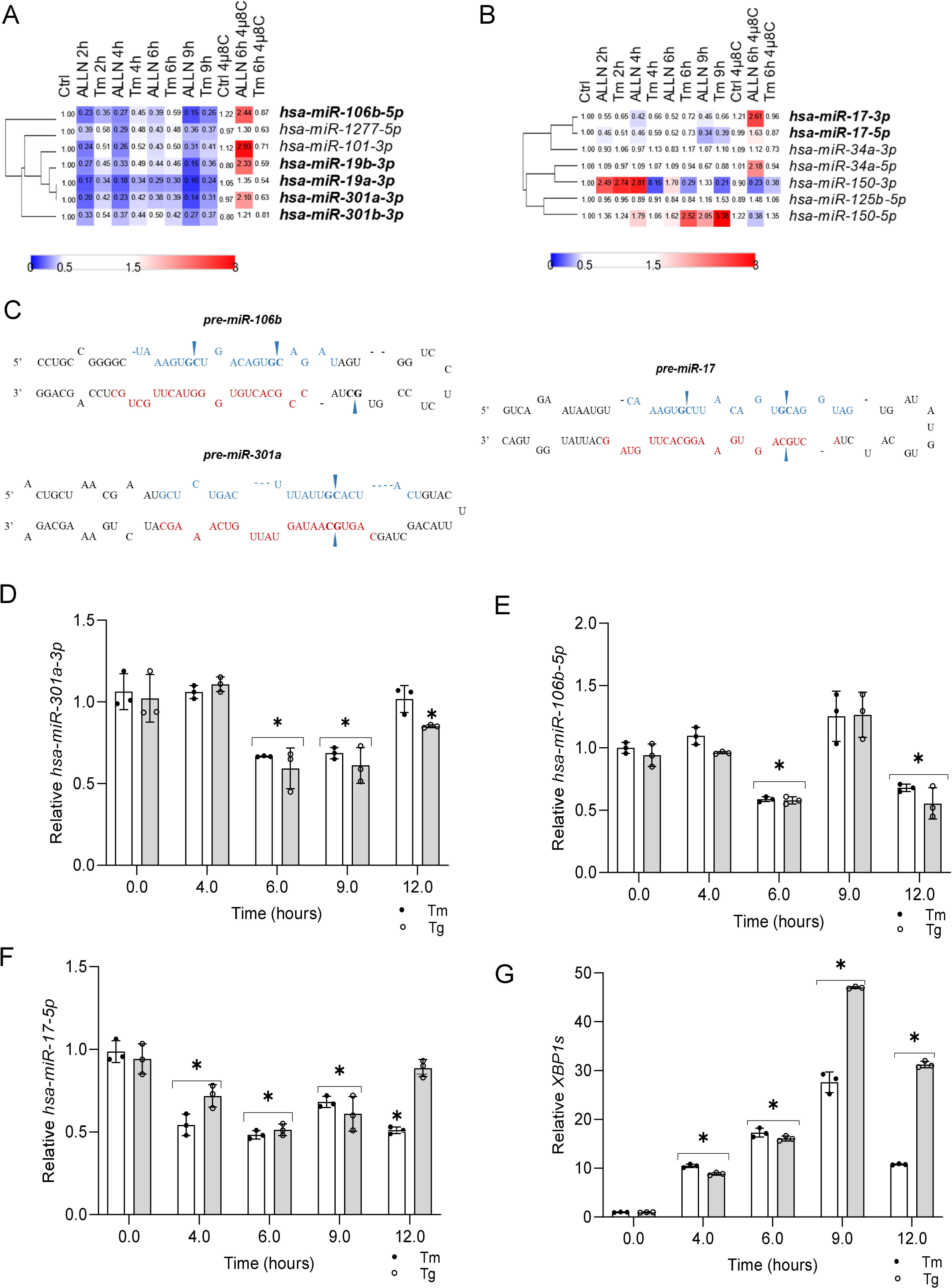
The impact of the IRE1 inhibition on ER-stress exposed 16HBE14o- genome-wide miRNA profiles. (**A**) miRNA expression was significantly (*P* < 0.05) restored upon IRE1 inhibition with 4µ8C in tunicamycin (Tm, 2.5 µg/ml) treated cells and ALLN – Calpain I inhibitor (100 µM) treated cells) based on NGS profiling. The 4µ8C was used at 20 µM concentration. Bold font indicates the presence of an IRE1 consensus motif. (**B**) Impact of IRE1 inhibition on miRNAs reported previously as IRE1 targets. Bold font indicates the presence of an IRE1 consensus motif. (**C**) Schematic representation of an IRE1 motif in selected miRNAs precursors (pre-miRNA) stem loops. Mature *miRNA-5p* sequences are marked with blue, and mature *miRNA-3p* sequences in red. Arrows mark the IRE1 consensus motif sequence. ER stress-induced changes in (**D**) *hsa-miR-301a-3p*, (**E**) *hsa-miR-106b-5p*, (**F**) *hsa-miR-17-5p*, and (**G**) *XBP1s* RNA levels in 16HBE14o- cells. The results from three independent experiments (*n* = 9) are plotted normalized to *RNU44* and *RPLP0* mRNA levels and expressed as a fold change over the no-stress controls. Error bars represent standard deviations. Significant changes (*P* value *P* < 0.05) are marked with an asterisk. ER stressors used: Tm (2.5 µg/ml), Tg (50 nM)).

As shown in **Figure 1A and B**, the initial selection was limited to 5 miRNAs that were reduced during ER stress and contained an IRE1 target sequence (bolded) as well as 2 miRNAs that were previously reported to be IRE1-dependent ones (42). Given the nonspecific effects of ALLN on IRE1 branch studies, we utilized a non-competitive inhibitor of the sarco/endoplasmic reticulum Ca ATPase, thapsigargin (Tg), as the second ER stress model for these studies. Consequently, based on the results of the independent, qPCR-based validation of the results from these two ER stress models (Tm and Tg), we selected *hsa-miR-301a-3p* and *hsa-miR-106b-5p* as potentially degraded by IRE1, and *hsa-miR-17-5p* as a previously identified target of IRE1 (42). Importantly, all three of the selected miRNAs contained IRE1 target sequence in their pre-miRNA sequences (**Figure 1C**). Next, we followed the selected miRNAs expression profiles during ER stress. As shown in **Figure 1D and E**, *hsa-miR-301a-3p* was reduced in cells exposed to either Tm or Tg for 6 and 9 hours, whereas reduction of *hsa-miR-106b-5p* was noted after 6- and 12-hours exposure. The putative control, *hsa-miR-17-5p,* was downregulated in both Tm and Tg treated cells after 4-, 6- and 9-hour exposure (**Figure 1F**). Notably, all tested miRNAs were reduced by about half and these reductions were negatively correlated with *XBP1s* transcript accumulation (**Figure 1G**) as a consequence of IRE1 activity. *XBP1s* mRNA began to accumulate after 4 hours of exposure to ER stressors and reached a maximum after 9 hours and remain elevated at 12 hours.

Inhibition of IRE1 activity with 4μ8C during ER stress not only prevented *XBP1s* mRNA and protein accumulation (**Figure 2A and B**), but it also restored the expression of *hsa-miR-17-5p*, *hsa-miR-301a-3p*, and *hsa-miR-106b-5p*, in a stressor independent manner (**Figure 2C, D, and E**). Furthermore, the expression of miRNA biogenesis components (*AGO2, DICER1 and DROSHA*) remained unaffected up to 12h of exposure to Tm-induced ER stress (**Figure 2F**), and that is in good agreement with previous reports (43). These data suggest that UPR-related reduction of these miRNAs is IRE1-dependent. Since the consensus sequence that targets selected miRNAs to the IRE1 endoribonuclease activity was present not only in mature selected miRNAs, but also in their respective pre-miRNAs, we tested if the precursors could be an IRE1 substrates. A complication is that pre-miRNAs are the result of pri-miRNA maturation and the pre-miRNAs are exported from nucleus to cytosol to be further processed to mature miRNAs (44). Furthermore, given that the nuclear envelope is continuous with ER membranes (45), IRE1 could process pre-miRNAs both in nucleus and cytosol. Hence, to test if IRE1 was degrading *pre-miR-17*, *pre-miR-301a* and *pre-miR-106b*, we compared levels of these precursors in nuclear and cytosolic fractions from cells exposed to ER stress with the pre-miRNAs obtained from cells treated with both ER stressors as well as the IRE1 inhibitor. *Pri-miR-17*, found only in the nucleus, was used as a quality control for the nuclear fraction, and mitochondrial encoded cytochrome b (*CYTB)* mRNA was used for the cytosolic control (**Figure 3A and B**).

**Figure 2.**
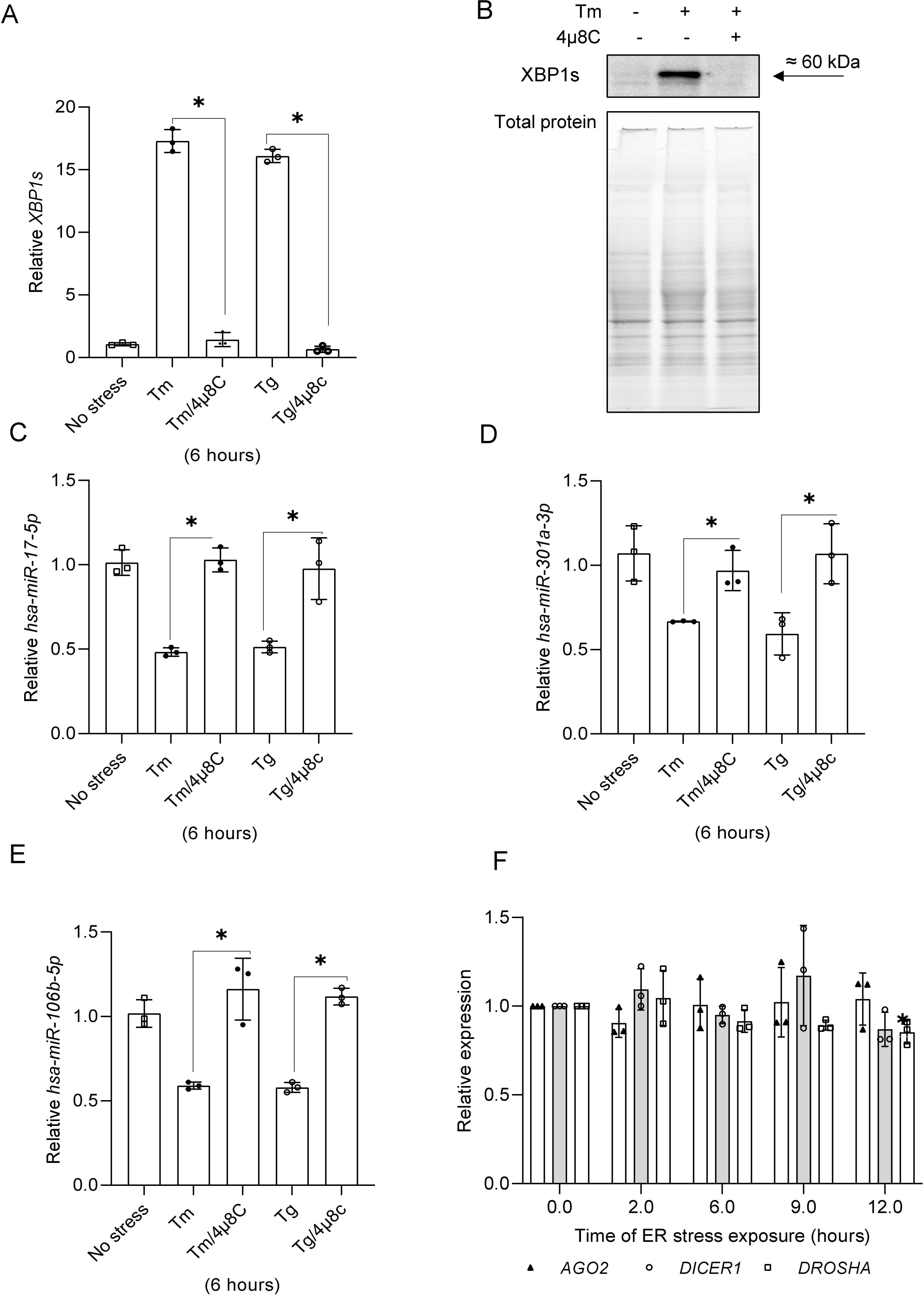
IRE1 inhibition during ER stress reduces XBP1s expression. **(A and B) and restores *hsa-miR-17-5p* (C), *hsa-miR-301a-3p* (D) and *hsa-106b-5p* levels (E).** hsa-miRs, and *XBP1s* mRNA levels were quantified with qRT-PCR and normalized to *RPLP0* mRNA levels, and expressed as a fold change over no-stress control samples. Data represents the mean ± SD of three independent experiments (3 replicates each). * P < 0.05 was considered significant. The changes in XBP1s protein levels were evaluated by western blots normalized to total protein levels. Changes in *hsa-miR-301a-3p*, *hsa-miR-106b-5p*, *hsa-miR-17-5p* were quantified by qRT-PCR and the results from three independent experiments (*n* = 9) were plotted normalized to *RNU44* levels and expressed as a fold change over the no-stress controls. Error bars represent standard deviations. Significant changes (*P* value *P* < 0.05) are marked with an asterisk. ER stressors used: Tm (2.5 µg/ml), Tg (50 nM)). The 4µ8C was used at a 20 µM concentration. (**F**) The Tm (2.5 µg/ml) treatment does not affect the miRNA biogenesis component expression. AGO2, DICER1 and DROSHA expression levels were obtained from 3 independent NGS-based analysis of transcriptome changes during ER stress (15,19). Error bars represent standard deviations.

**Figure 3.**
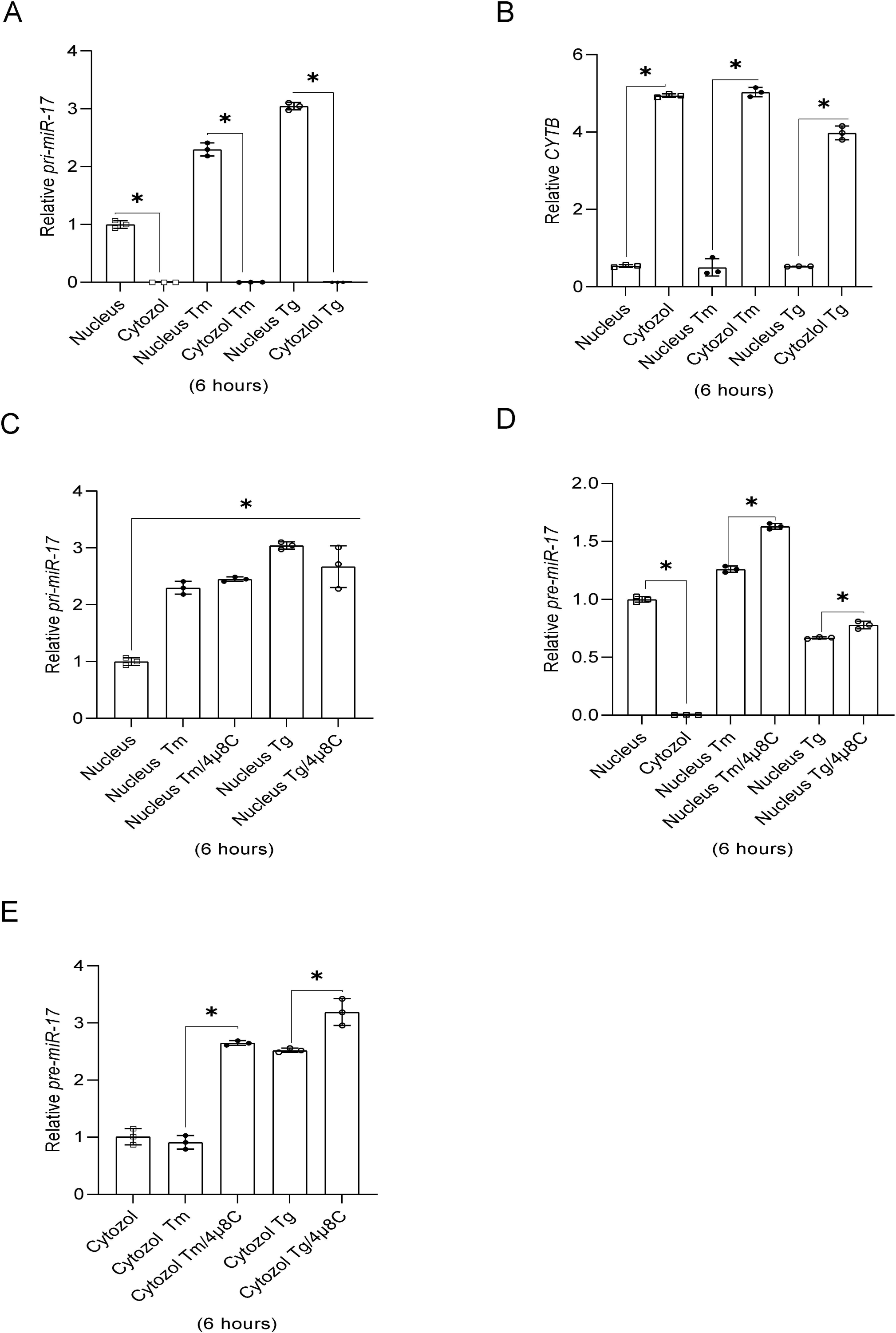
The pre-miR precursor of *hsa-miR-17-5p* is concentrated in the nuclear fraction and sensitive to IRE1 activity. qRT-PCR based analyses of nuclear and cytosolic fraction purities based on quantification of *pri-miR-17* (A) and *CYTB* (B) expression. RT-qPCR results from three independent experiments (*n* = 9) are plotted normalized to *RPLP0* levels and expressed as a fold change over the no-stress controls. Error bars represent standard deviations. Significant changes (*P* value *P* < 0.05) are marked with an asterisk. ER stressors used: Tm (2.5 µg/ml), Tg (50 nM)). The impact of 4µ8C (IRE1 inhibitor) on *pri-miR-17* (**C**) as well as *pre-miR-17* RNA levels in both nuclear and cytosolic fractions from ER stress exposed cells (**D** and **E**) were quantified with qRT-PCR and normalized to *RPLP0* (for *pri-miR-17*) or *RNU44* and expressed as a fold change over no-stress control samples. The results from three independent experiments (*n* = 9) are plotted. Error bars represent standard deviations. Significant changes (*P* value *P* < 0.05) are marked with an asterisk. ER stressors used: Tm (2.5 µg/ml), Tg (50 nM)). The 4µ8C was used at a 20 µM concentration.

The *pri-miR-17* was almost undetectable in the cytosolic fractions as expected. Despite the significant enrichment of *CYTB* in cytosol, much smaller amounts were also noted in nuclear fractions, presumably due to the presence of intimate contacts of mitochondria with nuclear envelope that have been proposed to be the gateway for mRNAs (46). As shown in **Figure 3C**, the expression of *pri-miR-17* was induced by ER stress and was not affected by the IRE1 inhibitor. In contrast, as shown in **Figure 3D and E**, *pre-miR-17* levels were significantly increased in both nuclear and cytosolic fractions from the cells treated with ER stressors in the presence of 4µ8C. Notably, the majority of *pre-miR-17* was detected in nuclear fractions (**Figure 3D**), which is in agreement with previous reports indicating that the majority of pre-miRNA is concentrated in the nucleus (47).

A similar pre-miRNA distribution was observed for both *pre-miR-301a* and *pre-miR-106b* that accumulated in the nucleus (**Figure 4**). Upon IRE1 inhibition, *pre-miR-301a* levels were significantly increased in both fractions, independent of ER stressor (**Figure 4A and B**). The *pre-miR-106b* levels were increased in nuclear fraction from the cells treated with Tm and Tg in the presence of 4µ8C (**Figure 4C**). This increase, however, was only found in the cytosolic fraction of Tm treated cells (**Figure 4D**). The lack of a 4µ8C effect in Tg treated cells in the cytosol could be a consequence of different ER stress models and the related UPR dynamics. Notably, as previously shown in **Figure 1E**, IRE1 inhibition increased mature *hsa-miR-106b-5p* levels in both Tm and Tg treated cells, so the lack of an effect for the pre-miR is unclear. To get better understanding of IRE1-mediated degradation of these miRs and pre-miRs, we compared their stability under normal conditions, and during ER stress with and without IRE1 inhibition (**Figure 5**). As shown in **Figure 5A and B**, both ER stress and IRE1 inhibition did not dramatically affect the mature hsa-miR-301a-3p nor hsa-miR-106b-5p. Although, in Tm-treated cells, the half-life of both of these miRNAs were about 1 hour shorter than in control conditions, and these molecules remained very stable with half-lives of about 10 hours, which is in good agreement with previous reports (48). Importantly, the stability of pre-miRs in the nucleus also remained unaffected (**Figure 5B and C**). In contrast, their half-life in cytosol was significantly reduced during the Tm treatment. The pre-miR-301a was reduced from 3.55 h to 2.43 h and pre-miR-106b was reduced from 3.05 h to 1.91h. Furthermore, the stabilities of pre-miRNA-106b and pre-miR-301a were completely restored to the values observed in the no stress conditions when IRE1 was inhibited (**Figure 5D and E**).

**Figure 4.**
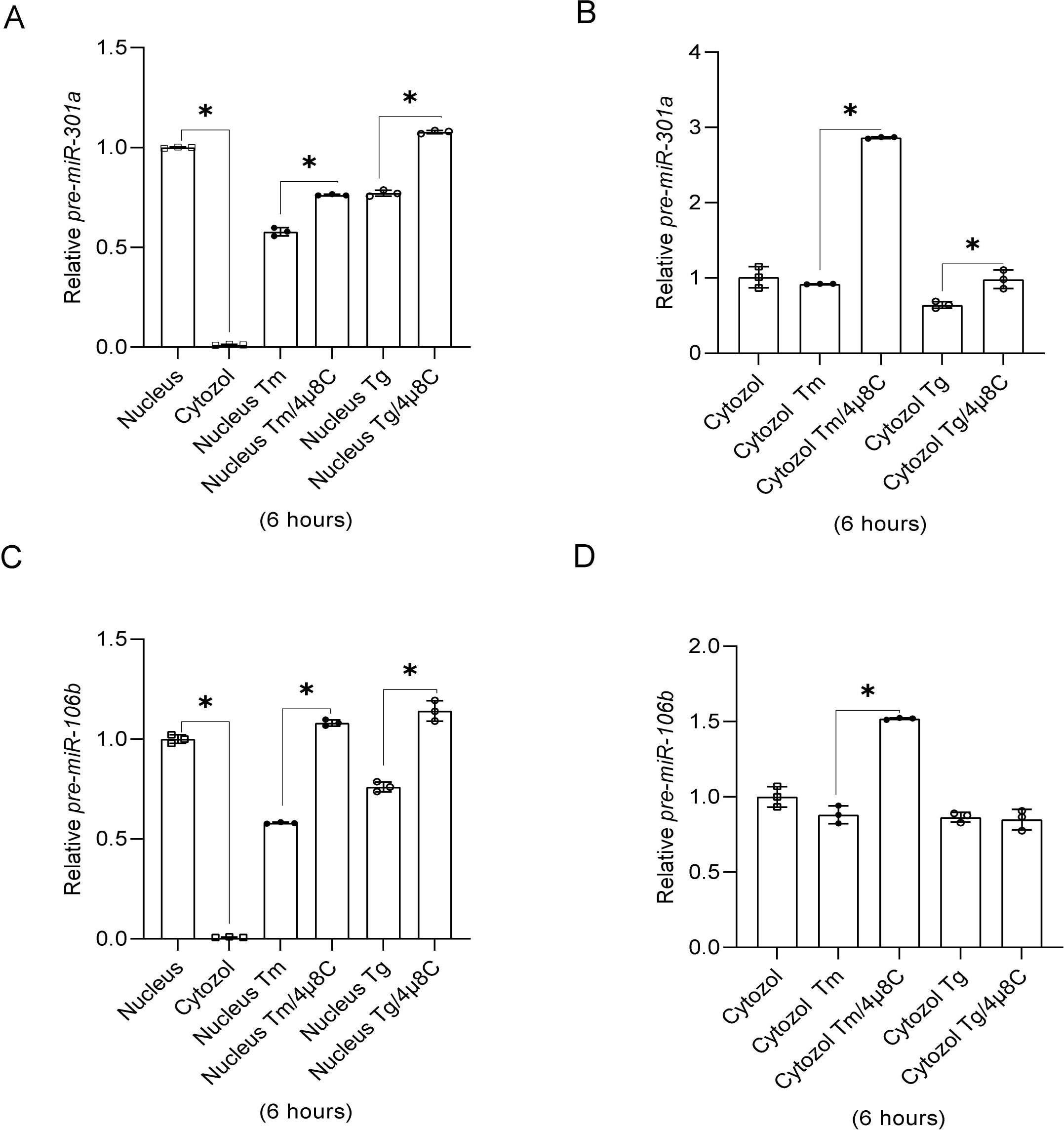
The pre-miR precursor of *hsa-miR-301a-3p* and *hsa-miR-106b-5p* are concentrated in nuclear fraction and sensitive to IRE1 activity. The impact of 4µ8C on *pre-miR-301a* (**A** and **B**) as well as *pre-miR-106b* in both nuclear and cytosolic fractions from ER stress exposed cells (**C** and **D**). RNA levels were quantified with qRT-PCR and normalized to *RNU44* and expressed as fold change over the no-stress control samples. The results from three independent experiments (*n* = 9) are plotted. Error bars represent standard deviations. Significant changes (*P* value *P* < 0.05) are marked with an asterisk. ER stressors used: Tm (2.5 µg/ml), Tg (50 nM)). The 4µ8C was used at a 20 µM concentration.

**Figure 5.**
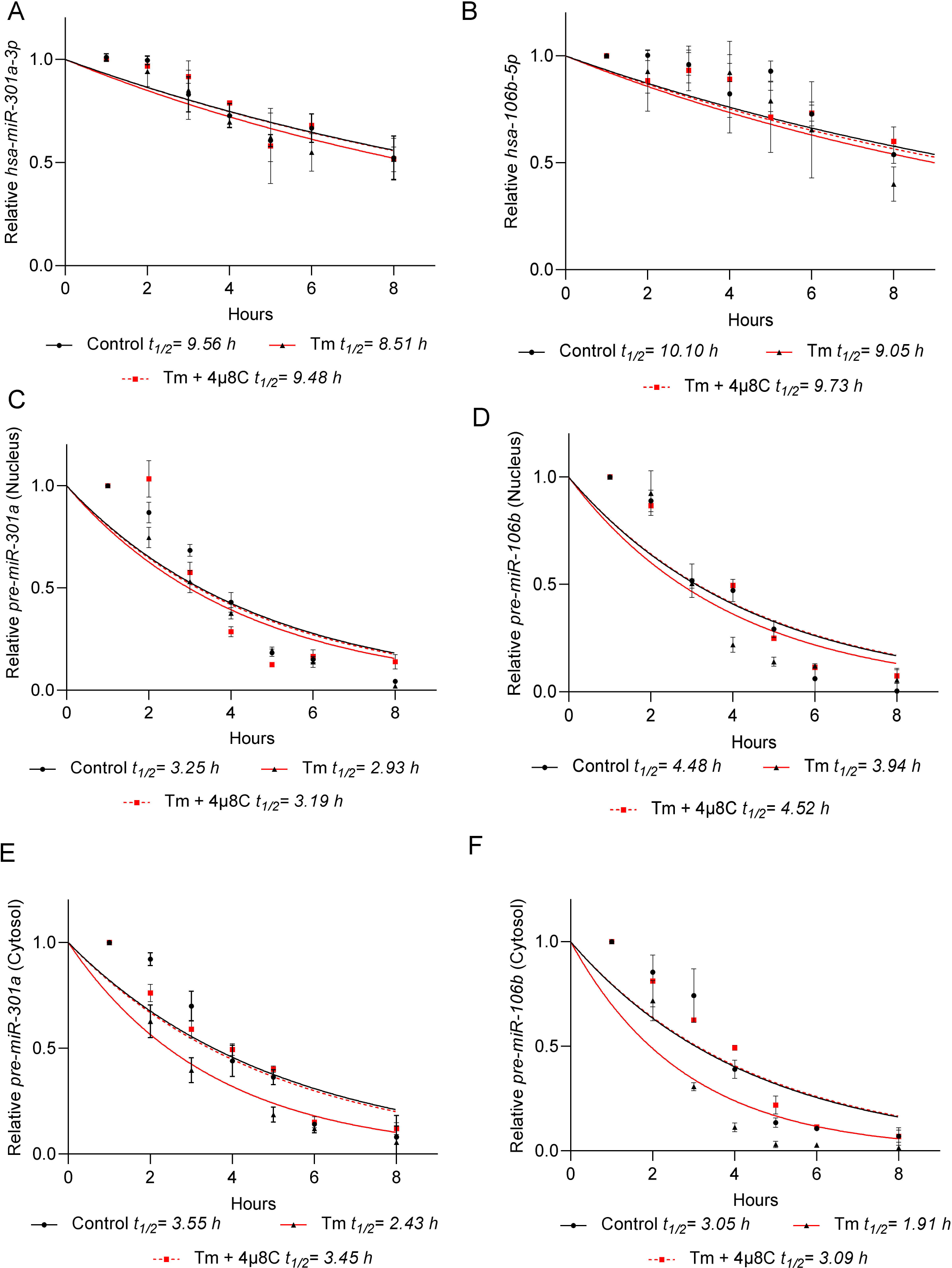
Inhibition of IRE1 activity during ER stress decreases *pre-miRNA stability in the cytosol*. RNA half-life measurements were taken in 16HBE14o- exposed to Tm (2.5µg/ml) in the presence or absence of 20 µM 4µ8C, and from the cells cultured in control conditions. Actinomycin D was added to stop transcription, after which the cells were collected, and total RNA was isolated from nuclear and cytosolic fractions. The levels of total *hsa-miR-301a-3p* (A), total *hsa-miR-106-5p* (B) nuclear pre-miR-301a (C), nuclear pre-miR-106b (D), cytosolic pre-miR-301a (E) and cytosolic pre-miR-106b (F) at each time point were measured by real-time PCR and normalized to endogenous *18S* rRNA levels. RNA values for each time point were calculated from 2 individual samples generated in at least 2 independent experiments and measured in 4 technical replicates. Relative RNA levels at the time points indicated were plotted as differences from RNA levels at the initial time point (*t* = 0). The mRNA half-lives were calculated from the exponential decay using the trend line equation *C*/*C*_0_ = *e*^−kdt^ (where *C* and *C*_0_ are RNA amounts at time *t* and at the *t*_0_, respectively, and *k*_d_ is the RNA decay constant). The error bars represent SD. The calculated parameters are provided in **Supplemental Table 1.**

At this point, the data supported the hypothesis that during UPR, IRE1 is reducing *hsa-miR-301a-3p* and *hsa-miR-106b-5p* via degradation of the respective pre-miRNAs in the cytosol, whereas the corresponding small differences in mature miRNA stability are the consequence of reduced pre-miRNA levels in cytosol. Although we block the pri-miRNA transcription, the pre-miRNA processing still remains active.

Next, we asked what the functional consequence of this IRE1-dependent regulation of miRNA expression during the UPR was. Our goal was to establish direct targets for both *hsa-miR-106b-5p* and hsa-*miR-301a-3p*. To do so, we analyzed our previous transcriptomic data obtained from 16HBE14o- cells exposed to Tm for 2-, 6- and 9-hours (15) in order to find transcripts that were significantly induced during ER stress and contained the conserved binding sited for these miRNAs. This approach resulted in selection of epiregulin (*EREG*) as a potential target of *hsa-miR-106b-5p*. Although this transcript was induced by ER stress (**Supplemental Figure 1B**), however, its levels remained unaffected when IRE1 was inhibited **(Supplemental Figure 1C**), and therefore we next moved on to *hsa-miR-301-3p*.

In this case, we focused on growth arrest and DNA damage inducible alpha (*GADD45A*) as a potential target of *hsa-miR-301a-3p*. The 3’ UTR region of *GADD45A* contains a 7-mer binding sequence for *miR-301a-3p* (**Figure 6A**). Notably, both the binding sequence and the miRNAs’ seed sequence are strongly conserved among different species (**Figure 6B and C**). *GADD45A* expression was induced by both Tm and Tg (**Figure 6D**), and IRE1 inhibition resulted in a significant reduction of this transcript levels (**Figure 6E**). Furthermore, when 4µ8C was added to 16HBE14o- cells exposed to Tm and Tg for 9 hours, when the maximum of *GADD45A* induction was observed, the levels of this transcript were dramatically reduced as well (**Supplemental Figure 1D**).

**Figure 6.**
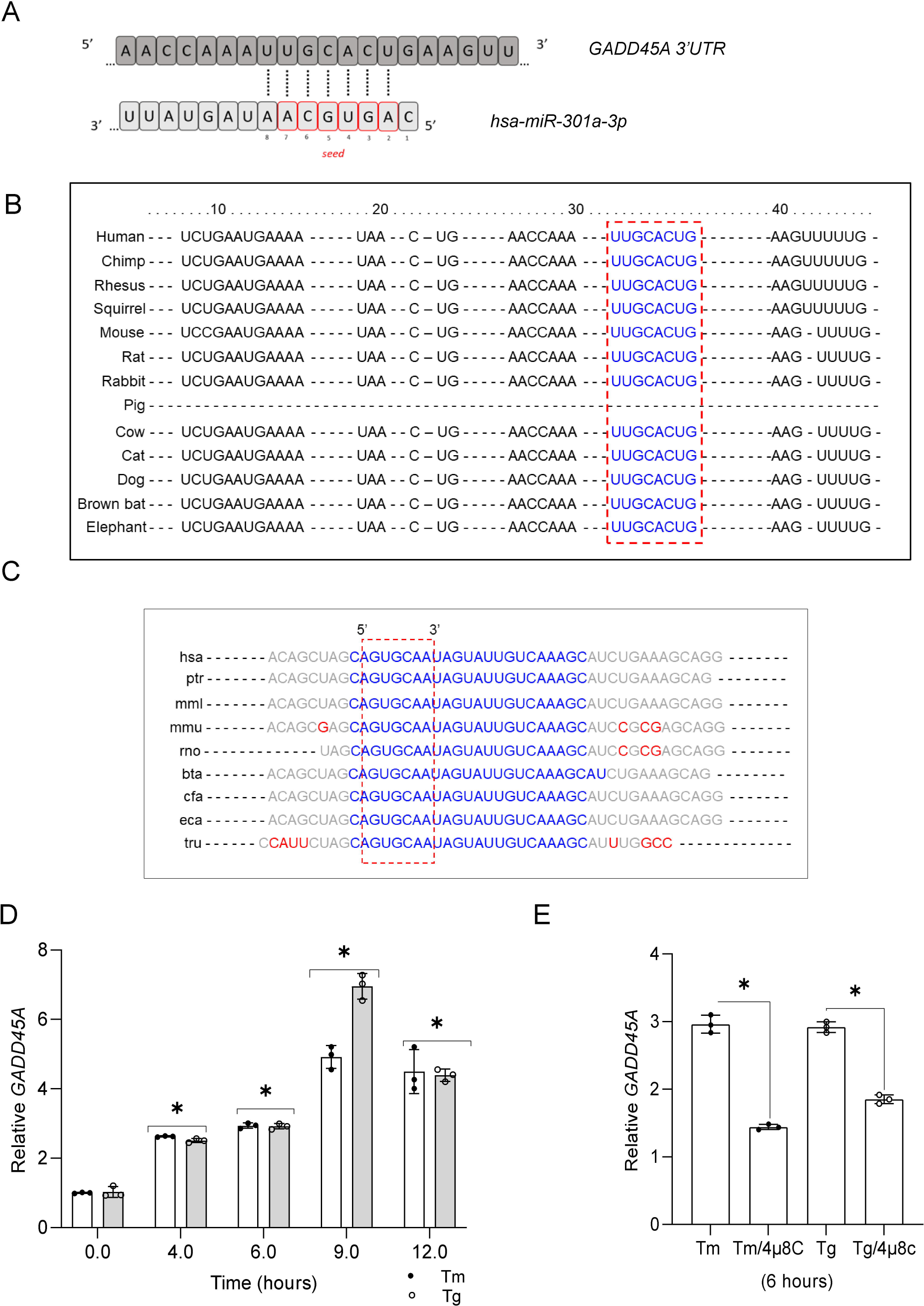
Inhibition of IRE1 activity during ER stress decreases *GADD45A* mRNA levels. Schematic representation of *hsa-miR-301a-3p* binding site in the *GADD45A* 3’ UTR sequence as predicted by Target scan 6.0 (**A**). The conservation of predicted binding site in *GADD45A* sequence, as marked with red box (**B**) and conservation of *hsa-miR-301a-3p* seed sequence were analyzed with Target scan, as marked with red box (**C**) (61). ER stress-induced changes in *GADD45A*, mRNA levels in 16HBE14o- cells (**D**) and the impact of 4µ8C on *GADD45A* expression (**E**) were analyzed with RT-qPCR and normalized to *RPLP0* mRNA levels, and expressed as a fold changeover no-stress control samples. The results from three independent experiments (*n* = 9) are plotted and expressed as a fold change over the no-stress controls. Error bars represent standard deviations. Significant changes (*P* value *P* < 0.05) are marked with an asterisk. ER stressors used: Tm (2.5 µg/ml), Tg (50 nM)).

Using a stable XBP1s-inducible cell line (20), we tested if *GADD45A* increased during the induction of *XBP1s* expression. As shown in **Figure 7A** and **B**, *GADD45A* expression was not affected by the increase in *XBP1s* mRNA. Next, we asked if restoring *hsa-miR-301a-3p* levels during ER stress with its precursor (premiR) affected *GADD45A* expression levels. We utilized *pre-miR-301a-3p* to limit the amount of mature miRNA and found that even under these conditions, the resulting *hsa-miR-301a-3p* levels were elevated by 400-fold in control conditions and by 200-fold during Tm treatment (**Figure 7C**).

**Figure 7.**
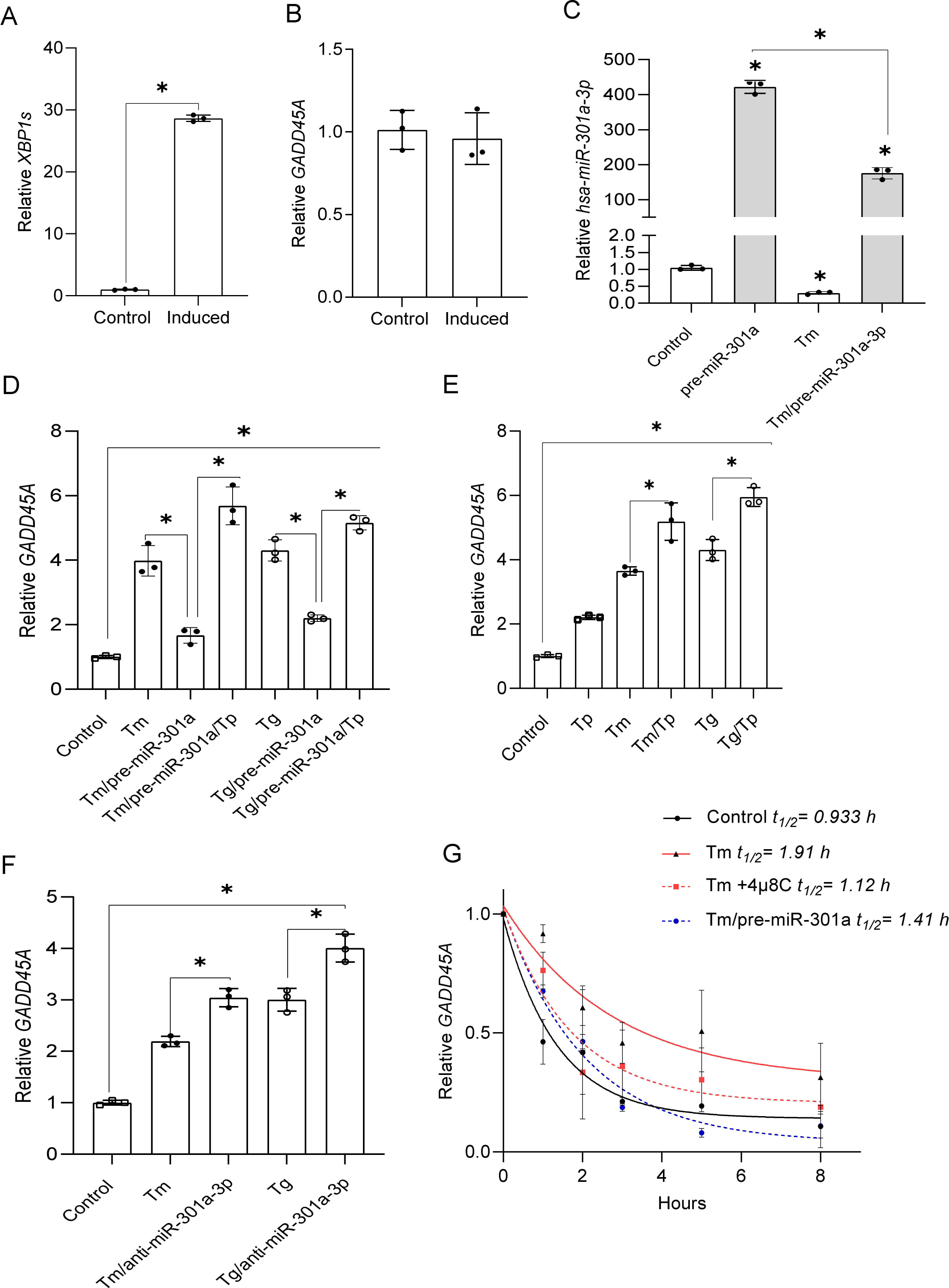
*Hsa-miR-301a-3p* regulates *GADD45A* expression and mRNA stability during ER stress in an XBP1s-independent manner. *XBP1s* expression was induced in the HeLa S3 cell line (20) and *XBP1s* (**A**) and *GADD45A* (**B**) mRNA levels were monitored with qRT-PCR and normalized to *RPLP0* mRNA levels, and expressed as the fold change over control (no induction) samples. Data represent the mean ± SD of three independent experiments (3 replicates each). * P < 0.05 was considered significant. 16HBE14o- cells were transfected with *pre-miR-301a* and specific target protector- Tp (**C D** and **E**) for the *GADD45A* binding site for *hsa-miR-301a-3p* or with anti-miR-301a-3p (**F**) or their respective controls and then treated with Tm (2.5 µg/ml) or Tg (50 nM) for 6 h. *GADD45A* mRNA levels were quantified by qRT-PCR and normalized to *RPLP0* mRNA. Data represent the mean ± SD of three independent experiments (3 replicates each). * P < 0.05 was considered significant. (**G**) *GADD45A* mRNA half-life measurements were taken in 16HBE14o- exposed to Tm (2.5µg/ml) in the presence or absence of 20 µM 4µ8C, as well as after pre-miR-301a transfection, and from the cells cultured under control conditions. Actinomycin D was added to stop transcription, after which the cells were collected, and total RNA was isolated and *GADD45A* mRNA levels at each time point were measured by real-time PCR and normalized to endogenous *18S* rRNA levels. RNA values for each time point were calculated from 2 individual samples generated in at least 2 independent experiments and measured in 4 technical replicates. Relative RNA levels at the time points indicated were plotted as differences from RNA levels at the initial time point (*t* = 0). The mRNA half-lives were calculated from the exponential decay using the trend line equation *C*/*C*_0_ = *e*^−kdt^ (where *C* and *C*_0_ are mRNA amounts at time *t* and at the *t*_0_, respectively, and *k*_d_ is the mRNA decay constant). The error bars represent SD. The calculated parameters are provided in **Supplemental Table 1**.

To demonstrate a direct interaction between *miR-301a-3p* and *GADD45A* mRNA, we used a specific target protector (Tp, target mask) molecule (49). The target protector is a morpholino uniquely complementary to *hsa-miR-301a-3p* target sequence in the *GADD45A* 3’UTR, and interferes with a single miRNA-mRNA pair by binding specifically to the miRNA target sequence in the 3′ UTR (49). As shown in **Figure 7D**, overexpression of *pre-miR-301a* in 16HBE14o- cells resulted in a significant reduction of *GADD45A* mRNA in both Tm and Tg treated cells. Furthermore, the Tp reversed the inhibitory effect of *hsa-miR-301a-3p* on *GADD45A* mRNA levels in ER stressed cells. The Tp was also effective in elevating *GADD45A* expression in the absence of *pre-miR-301a* transfection, in both unstressed and Tm and Tg treated cells (**Figure 7E**). Since this data suggested that despite the IRE1-dependent degradation, the remaining levels of *hsa-miR-301a-3p* were still effectively modulating or buffering *GADD45A* expression, we tested if antagomiR (anti-miR-301a-3p) will allow further increase of this transcript in ER stressed cells. As shown in **Figure 7F**, antagomiR transfection resulted in elevated *GADD45A* mRNA levels in both Tm and Tg treated cells, lending further support to the regulation of *GADD45A* mRNA by *miR-301a-3p.* Finally, we have measured *GADD45A* mRNA stability in no stress conditions and during Tm induced ER stress in the presence of IRE1 inhibitor or with the addition of pre-miR-301a (**Figure 7G**). The *GADD45A* mRNA was stabilized during the UPR and the mRNA half-life increased from about 1 h to 1.91 h. IRE1 inhibition restored this transcript stability to control levels (half-life of 1.1 h) during Tm treatment. These results are in good agreement with the previous report of a 45 minute *GADD45A* mRNA half-life (50). Furthermore, significant *GADD45A* mRNA destabilization was also observed when the cells were transfected with *pre-miR-301a* (half-life of 1.4 h) during Tm treatment.

Next, we tested if IRE1 inhibition also resulted in increased GADD45α protein levels in Tm treated cells. As shown in **Figure 8A and B**, consistent with our mRNA results, GADD45α protein levels in the presence of 4µ8C were significantly decreased. Furthermore, increasing *hsa-miR-301a-3p* levels with its pre-miR resulted in a dramatic reduction of GADD45α expression in Tm treated cells as well (**Figure 8C and D**). A similar negative effect of *hsa-miR-301a-3p* on GADD45α was also observed when cells were exposed to Tg (**Supplemental Figure 2AB**). Furthermore, silencing *ERN1* during ER stress (51) not only led to the reduction of *XBP1s* mRNA levels, but also resulted in increased expression of all tested miRNAs (*hsa-miR-17-5p*, *hsa-miR-301a-3p* and *hsa-miR-106b-5p*) and reduced *GADD45A* mRNA levels (**Supplemental Figure 3**).

**Figure 8.**
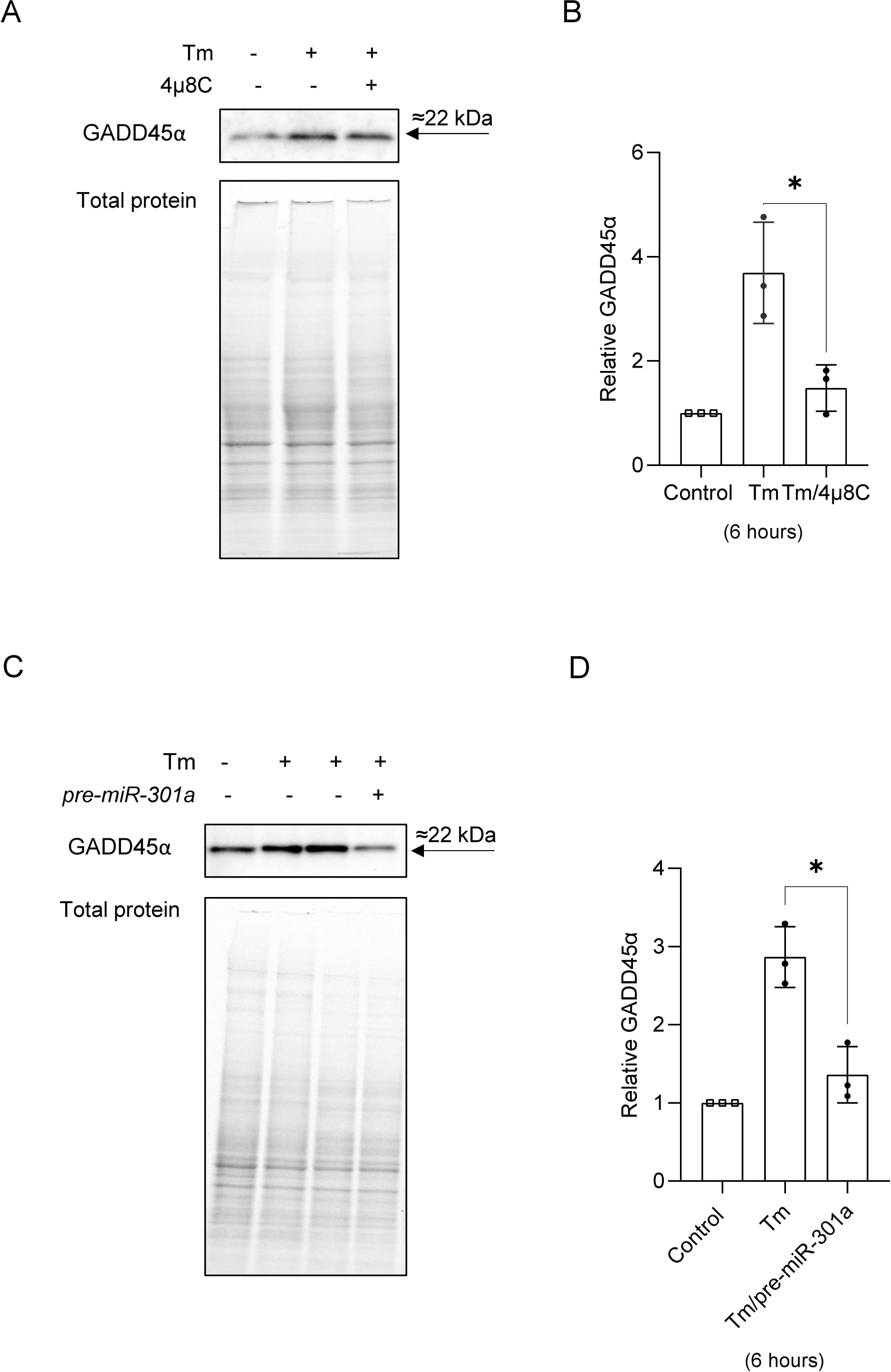
IRE1 and *pre-miR-301a* affect GADD45α levels during ER stress. 16HBE14o- cells treated with and without an IRE1 inhibitor (4µ8C) and were treated with Tm (2.5 µg/ml) for 6 h (**A and B**). The 16HBE14o- cells were first transfected with *pre-miR-301a-3p* or scramble control and two days later were treated with Tm (2.5 µg/ml) for 6 h (**C** and **D**). The corresponding changes in GADD45α protein levels were monitored with Western blot that was normalized to total protein levels (**A** and **C**), and related to the respective control. Data represent the mean ± SD of 3 independent experiments. *P < 0.05 was considered significant.

Taken together, the data clearly demonstrates that *hsa-miR-301a-3p* affects not only *GADD45A* mRNA expression, but importantly, also GADD45α protein expression during ER stress.

This data suggested that IRE1’s reduction of *hsa-miR-301a-3p* levels and the corresponding increases *GADD45A* expression could contribute to cell death decisions during ER stress. To test this, we performed real time and label free holographic microscopy-based monitoring of cell death and viability using a HoloMonitor**®** time-lapse cytometer. Holographic microscopy was used to follow the optical thickness and irregularity of cells exposed for up to 24 hours to Tm in the presence or absence of *pre-miR-301a* (**Figure 9A**). Healthy cells are irregular in shape and thin, whereas dying cells are round and thick (30–35). As shown in **Figure 9B and C**, a slight trending of dying cells was observed starting after 12 hours exposure to Tm which significantly increased by 24 hours and these changes correlated well with the percentage of healthy cells that is drastically declining after 24 hours of Tm treatment. In contrast, Tm treated cells in the presence of *pre-miR-301a* remained healthy up to 24 hours with no significant difference in the percentage of dying and healthy cells compared to the no stress controls (**Figure 9C**).

**Figure 9.**
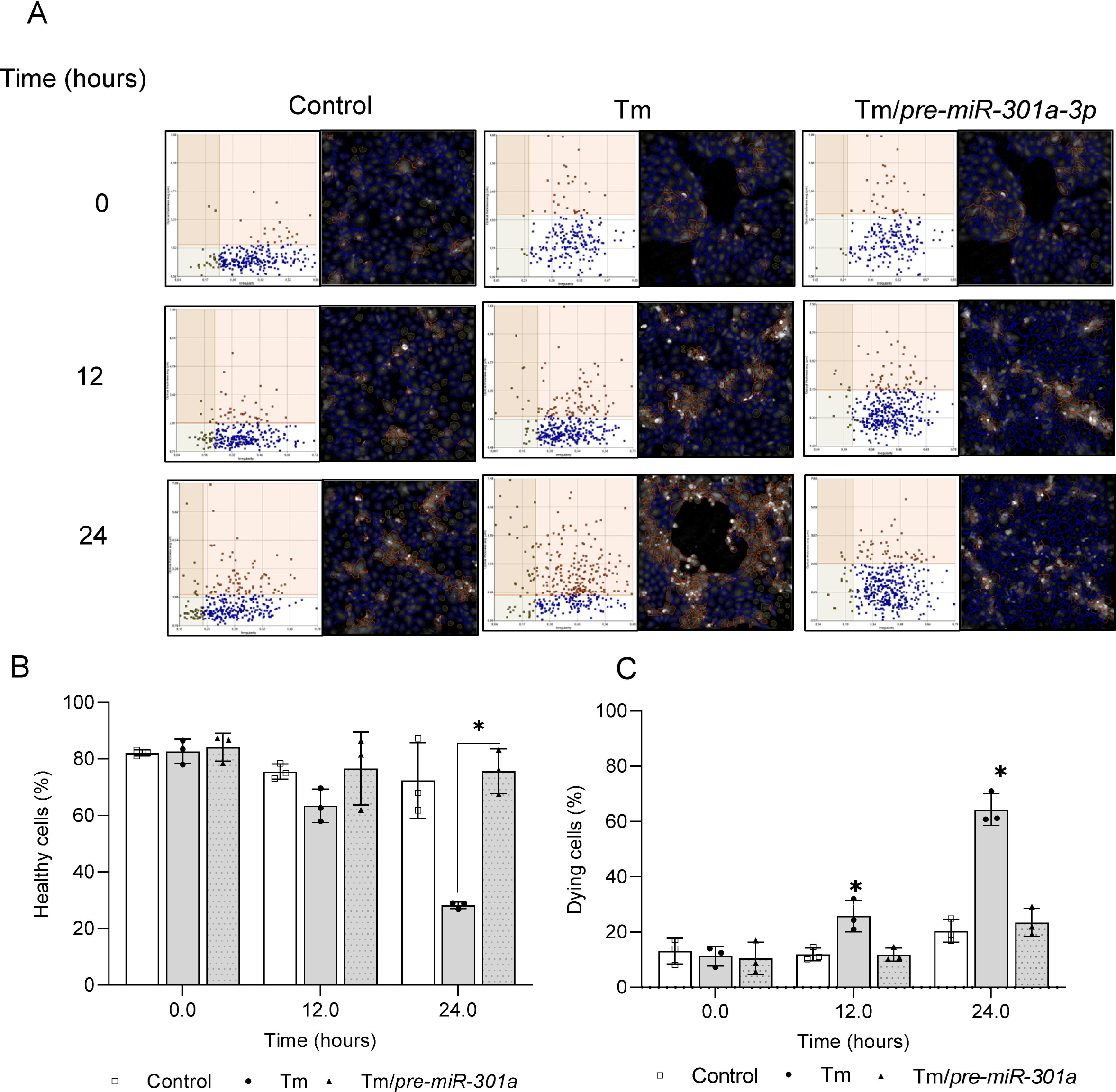
IRE1-mediated degradation of *pre-miR-301a* during ER stress contributes to cell death decisions. The results of real-time monitoring of cell viability with the real time and label free holographic microscopy using a HoloMonitor M4**®** time-lapse cytometer of 16HEB14o- cells transfected with *pre-miR-301a-3p* or the scramble control and 48 hours later treated with Tm (2.5 µg/ml) up to 24 h. Images were collected every 15 minutes (from 5 independent optical fields), and the distribution of live (blue) and dying cells (red) as well as dead cells (grey) based on their optical thickness and irregularity is presented at the 12 and 24 hour time points. The images from up to 5 independent optical fields were collected and analyzed according to manufacture instructions with HoloMonitor® App Suite software. Representative samples are shown (**A**). For all analyses, the same cell parameters qualifications were applied. Experiments were performed in triplicate. Based on the cells irregularity and average optical thickness the percentages of healthy cells (**B**) and of dying cells (**C**) were calculated. Data represent the mean ± SD of three independent experiments. * P < 0.05 was considered significant. Schematic representation of role of IRE1-mediated degradation of *pre-miR-301a* on cell fate decision during ER stress (**D**).

To confirm that the interaction of *hsa-miR-301a-3*p with the binding site in *GADD45A* 3’UTR contributes to cell fate decisions, we used the same approach to follow the 16HBE14o- viability during ER stress in the presence of the specific target protector. As shown in **Figure 10**, these experiments confirmed that specific protection of *GADD45A* mRNA from *hsa-miR-301a-3p* mediated degradation with the target protector significantly accelerated the cell death in Tm treated 16HBE14o-. Although the transfection with the Tp control was increasing cell death in both control cells and Tm treated in the ER stress exposed cells, the percentage of healthy cells was significantly lower in the presence of Tp starting from 6 hours (**Figure 10B**) that was well reflected in the percentage of dying cells (**Figure 10C**).

**Figure 10.**
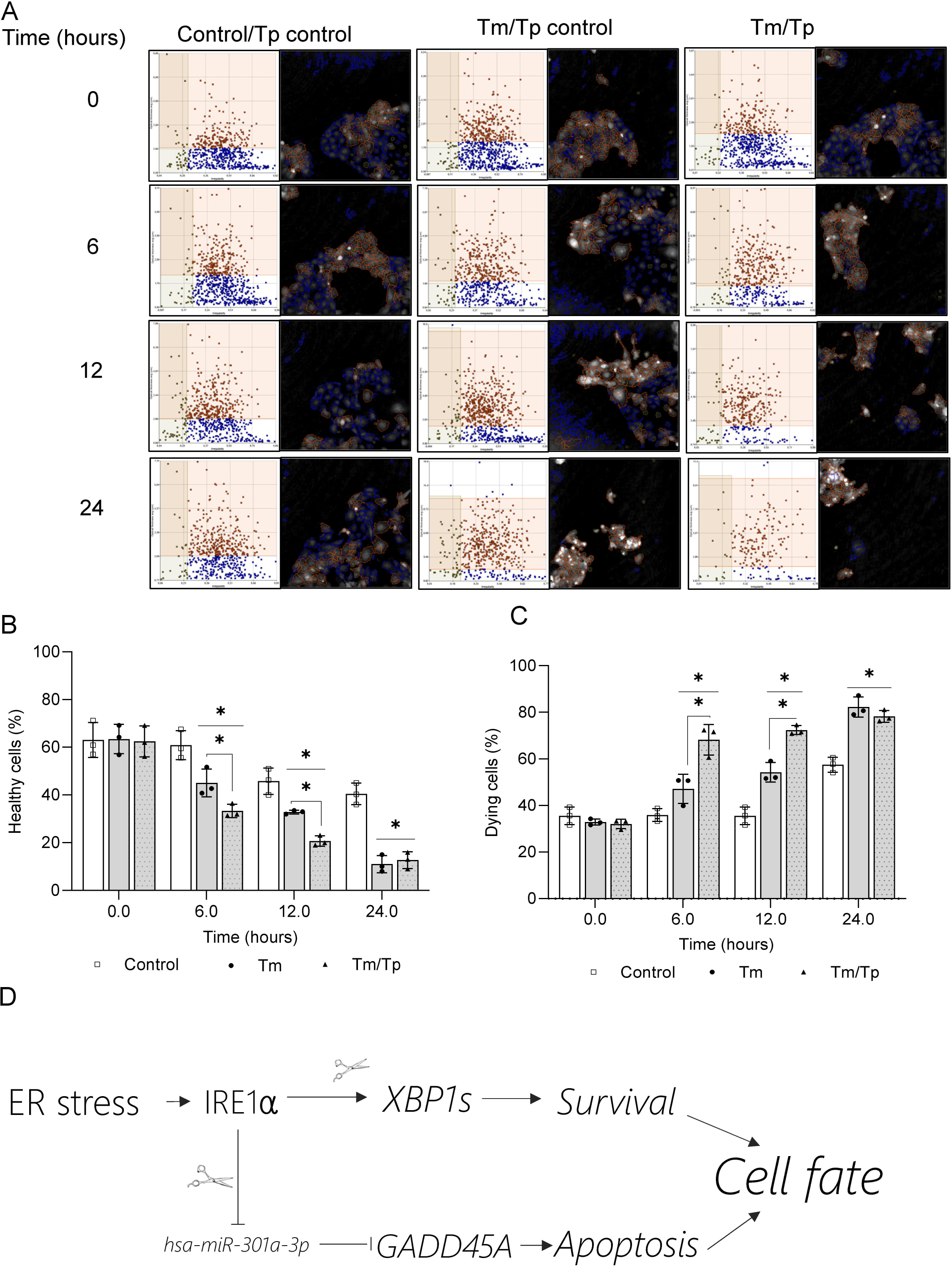
*hsa-miR-301a-3p* modulates *GADD45A* levels during ER stress and contributes to cell death decisions. The results of real-time monitoring of cell viability using holographic microscopy (HoloMonitor M4**®** time-lapse cytometer). 16HBE14o- cells were transfected with target protector (Tp) or Tp control and treated with Tm (2.5 µg/ml) up to 24 h. Images were collected every hour from 5 independent optical fields. The distribution of live (blue), dying cells (red), and dead cells (grey) were based on their optical thickness and irregularity is presented only at the 6- and 12-hour time points (**A**). The images from up to 5 independent optical fields were collected and analyzed according to manufacture instructions with HoloMonitor® App Suite software. One representative optical field is presented in the right panels. For all of the analyses, the same cells parameter quantification was applied as in Figure 8. Experiments were performed in triplicates. Based on the cells irregularity and average optical thickness the percentages of healthy cells (**B**) and of dying cells (**C**) were calculated. Data represents the mean ± SD of three independent experiments. * P < 0.05 was considered significant. Schematic representation of role of IRE1-mediated degradation of pre-miR-301a on the cell fate decision during ER stress (**D**).

Taken together, our data indicate that during UPR, IRE1 degrades *pre-miR-301a* and thus enhances the expression of *GADD45A* mRNA and protein which promotes the ER stressed induced cell apoptosis (**Figure 10D**). This illustrates another function of IRE1 during the UPR.

## Discussion

The UPR functions to restore ER homeostasis or in the presence of unmitigated stress conditions to promote cell death. In the present study using immortalized human bronchial epithelial cells and pharmaceutical stressors, we focused on the role of IRE1, the most evolutionarily conserved UPR signaling pathway, to understand IRE1’s potential role in the demarcation between the pro-survival programs and apoptosis. IRE1 is known to utilize its endoribonuclease activity for 2 processes: 1) to splice *XBP1* mRNA to generate a functional transcription factor to induce a pro-survival pathway and 2) to cleave and inactivate mRNAs and miRNAs to reduce the ER load (9,14,41,52–55). IRE1 also induces the stress pathways driven by JUN N-terminal kinase (JNK) and by nuclear factor-κB (NF-κB) (reviewed in: (4)). The goal here was to identify potential miRNAs that would influence either survival or apoptosis.

Previous reports including our own have reported that the global miRNA levels during ER stress are reduced during the UPR adaptive response (19,56). Given that the that IRE1-mediated RNA degradation involves miRNA (42,52), it was plausible that this enzyme might be more directly responsible for UPR-related miRNA reduction in addition to the obvious loss of miRNAs in the RISCs. In our previous research, when we followed genome wide expression changes in ER stressed immortalized human bronchial epithelial cells, 16HBE14o- (19), only 7 of the miRNAs were significantly reduced by at least 2 fold independent of the ER stress model in all the times tested (2-, 4-, 6- and 9-hours). 16HBE14o- cells are a commonly used model for studying human airway epithelia functions under both physiological and pathological conditions (18,57). This approach, however, did not consider the possibility that IRE1’s negative effects on miRNAs levels could be masked by the induction of these ncRNAs by the PERK and ATF6 UPR branches.

Previous studies had indicated that an important role of IRE1 in regulating miRNA levels during UPR that includes both maturation and degradation of these noncoding RNAs (52–55). In mouse models, IRE1 has been shown to be responsible for a Dicer-independent maturation of pre-miR-2317 (54), as well as responsible for degradation of group of miRNAs (*miRs-17, −34a, −96, −125b*) that normally repress translation of *Caspase-2* mRNA (53). In mouse fibroblasts, inhibition of IRE1 resulted in accumulation of miR-150 (55). The functional involvement of IRE1 in UPR-related regulation of human miRNAs, however, has not been reported. Our results directly support IRE1’s ability to cleave pre-miRNAs such as was shown for pre-miR-17 [16].

We used two pharmacological stress conditions to activate the UPR and also pretreat cells with or without an IRE1 inhibitor (4μ8C) that inhibits IRE1’s endonuclease activity (39) in order to isolate the effect of IRE1 in decreasing miRNAs levels. From Next-Generation sequencing analyses, we identified 12 miRNAs that were downregulated during tunicamycin treatment. During ER stress conditions, we treated the cells with or without the IRE1 endonuclease inhibitor and found that 3 of these miRNAs, *miR-301a-3p*, *miR-106b-5p*, and *miR-17-5p* were elevated after IRE1 endonuclease inhibition, suggesting the IRE1 was degrading these miRNAs. As a further test that the inhibitor was working under these conditions, we demonstrated that spliced *XBP1* mRNA levels were also dramatically reduced by this treatment. Furthermore, one of these three microRNAs, mouse *pre-miR-17*, has been previously shown to be directly cleaved by recombinant human IRE1 in T-REx-293 cells at 3 cleavage sites including in a mature mouse *miR-17-3p* site (41).

We used pre-miRNA for two reasons. First, previous studies had indicated that IRE1 could promote maturation or degradation of them as mentioned above. Two, we wanted to limit the number of mature miRNAs in our transfections in order to limit the non-specific changes in gene expression that can sometimes occur with supraphysiological levels of mature miRNAs (58). In this case, the use of pre-RNAs increased the miRNA levels 400-fold compared to the use of miRNA mimics which can elevate the cellular levels of miRNAs to hundreds of thousands of times (24,59,60).

To confirm these effects in airway epithelial cells, we tested IRE1’s role in processing or degrading *pri-miR-17* or *pre-miR-17* and whether these potential processes occurred in the nucleus or cytosol after 6 hours of tunicamycin or thapsigargin treatment with or without the IRE1 inhibitor 4μ8C. The results indicated the *pre-miR-17*, but not *pri-miR-17*, was decreased during both stressor treatments and this decrease was reversed in the presence of 4μ8C. Furthermore, the results indicated that this occurred in both the nucleus and the cytosol. This supports previous studies by Scott Oakes and colleagues that in cell-free systems that recombinant IRE1α endonucleolytically cleaved mouse *pre-miR-17* precursor at sites distinct from DICER, but did not affect the *pri-miR-17* levels (41). Our results in human airway epithelial cells confirm their cell-free analyses.

Using the same analysis, we then tested the other two immature miRNAs, *pre-miR-301a* and *pre-miR-106b*. For *pre-miR-301*, interestingly, most of the pre-miR is in the nucleus and very little in the cytosol. Furthermore, in both the nucleus and the cytosol, IRE1 inhibition in the presence of both stressors significantly elevates their expression, suggesting that IRE1 endonuclease activity is decreasing their levels during stress responses just like *pre-mir-17*. For *pre-miR-106b*, the same effect was seen with both stressors and the IRE1 inhibitor in the nucleus, but interestingly, it was only seen during tunicamycin treatment in the cytosol, but not seen with thapsigargin treatment and inhibitor. However, our follow up analyses of miRNA and pre-miRNA stability clearly indicate that in the ER stress-exposed cells, IRE1 is processing cytosolic pre-miRNAs, whereas the changes in the nuclear fraction of pre-miRNA are a consequence of the IRE1-containnig ER membrane continuity with the nuclear envelope (**Supplemental Figure 2C**).

The potential targets of *miR-301a-3p* included an exact match for the 3’ UTR of *GADD45A,* a known proapoptotic gene. Furthermore, this miRNA targeting sequence is well-conserved in a number of species. We monitored the activity of *GADD45A* mRNA levels during both stress conditions and found that IRE1 inhibition significantly decreased the *GADD45A* mRNA levels indicating that the endonuclease activity of IRE1 was either influencing *GADD45A* mRNA directly or indirectly through spliced XBP1 or a miRNA that affected *GADD45A* mRNA levels such as *miR-301a-3p*. To test IRE1’s effect was occurring through XBP1s, we used an inducible *XBP1s* cell line, and first tested if induction of *XBP1s* had any influence on *GADD45A* mRNA and found that *GADD45A* levels did not change.

To test the role of *miR-301a-3p*, we utilized a target protector for the putative miR-301a-3p binding site in the 3’UTR of *GADD45A*. Using 16HBE14o- cells that were induced with either tunicamycin or thapsigargin, we found that both methods increased *GADD41A* levels, and that *pre-miR-301* transfection significantly decreased *GADD45A* levels in both stressors, and importantly, that the addition of a target protector to the *miR-301a-3p* binding site in *GADD45A* 3’UTR was able to reverse this effect and increase *GADD45A* levels. Further support came from the use of an antagomir (*anti-miR-301a-3p*) that enhanced the *GADD45A* mRNA levels as well. Together, this indicated that IRE1’s inhibition of a *pre-miR-301a* during stress conditions facilitated *GADD45A* mRNA level increases that could lead to a pro-apoptotic response. It also indicated that the IRE1 activity could influence both pro-survival (splicing *XBP1*) and pro-apoptotic signaling pathways by degrading *pre-miR-301a* to elevate *GADD45A* levels. To confirm this hypothesis, we tested if during ER stress that inhibition of IRE1 endonuclease activity would (1) decrease *GADD45A* mRNA levels and increase *pre-miR-301* levels and if this would (2) decrease GADD45α protein and both of these were confirmed.

IRE1’s effect suggested that its endonuclease activity prevented the maturation of a miRNA that targets a pro-apoptotic gene, *GADD45A,* and therefore enhanced *GADD45A* expression during stress conditions. The other better-known effect of IRE1 is its endonuclease activity on unspliced *XBP1* mRNA to generate spliced *XBP1s* mRNA that leads to a functional transcription factor that promotes cell survival. To test this model and the role of *miR-301a-3p* in this process, we performed real time and label free holographic microscopy-based monitoring of cell death and viability using HoloMonitor**®** time-lapse cytometer (30–35). We followed the optical thickness and irregularity of cells exposed for up to 24 hours to Tm in the presence or absence of *pre-miR-301a*. This analysis indicated that under control conditions greater than ∼10% of the cells are thick and round (dying), and after 12 and 24 hours of tunicamycin treatment this increased to ∼25 and ∼65% were dying. After transfection with *pre-miR-301a*, however, these numbers decreased to ∼10% and ∼25%, respectively, indicating that this miRNA dramatically reversed the level of apoptosis. This supports the idea that the IRE1 pathway has components that both support survival as well as apoptosis, and the balance of these responses and the level of *GADD45A* dictates the final result. A schematic model of this overview is shown in **Figure 10D**.

To confirm that the interaction of *hsa-miR-301a-3p* with the binding site in *GADD45A* 3’UTR contributes to cell fate decisions, we used the same approach to follow the 16HBE14o- viability during ER stress in the presence of the specific target protector. These experiments confirmed that specific protection of *GADD45A* from *hsa-miR-301a-3p*-mediated degradation with the target protector significantly accelerated the cell death in Tm treated 16HBE14o-. Although the transfection with the control Tp did increase cell death levels in both control cells and Tm treated, in the ER stress exposed cells, the percentage of healthy cells was significantly lower in the presence of *GADD45A* 3’ UTR Tp starting at 6 hours. This was also well reflected in the percentage of dying cells. Taken together, the data indicate that during UPR, IRE1 degrades *pre-miR-301a* and thus enhances the expression of *GADD45A* and this switches the ER stressed cells to apoptosis. In summary, our analysis indicated that IRE1’s endonuclease activity is a two-edged sword that can both promote survival through spliced increased *XBP1* expression or cell death through degradation of a miRNA that inhibits *GADD45A* expression levels during the UPR.

## Abbreviations

4µ8C: IRE1 Inhibitor III
16HBE14o: immortalized human bronchial epithelial cells
ATF6: activating transcription factor 6
ALLN: inhibitor of calpain 1
BIP: binding immunoglobulin protein
BSA: bovine serum albumin
id: identification
CYTB: Cytochrome B
DMSO: dimethyl sulfoxide
ER: endoplasmic reticulum
EREG: epiregulin
FBS: fetal bovine serum
GADD45A: growth arrest and DNA-damage-inducible alpha
GAPDH: glyceraldehyde-3-phosphate dehydrogenase
HeLa: human cervix adenocarcinoma cells
IRE-1: inositol-requiring enzyme 1
miRNA: microRNA
NGS: next generation sequencing
PERK: PKR-like endoplasmic reticulum kinase
PBS: phosphate buffer saline
qRT-PCR: quantitative real-time PCR
RIDD: IRE-1 dependent decay
RNU6: U6 small nuclear
RNU44: U44 small nuclear
RPLP0: ribosomal protein lateral stalk subunit p0
SD: standard deviation
SDS: sodium dodecyl sulphate
SDS-PAGE: sodium dodecyl sulphate–polyacrylamide gel electrophoresis
Tm: tunicamycin
Tg: thapsigargin
UPR: unfolded protein response
UTR: untranslated region
XBP-1s: spliced X-box binding protein 1

## Declarations

### Ethics statement

Not applicable

### Competing Interest

The authors declare no conflict of interest.

### Funding

This research was funded by National Science Center “Opus” Program under contract UMO-2020/37/B/NZ3/00861 (R.B.).

### Data Availability

All data generated or analyzed during this study are included in this published article (and its supplementary information files). Deep sequencing data were deposited in Gene Expression Omnibus (GEO) at accession numbers: GSE117629 and GSE129813

### Author Contributions

Magda Gebert Sylwia Bartoszewska and Lukasz Opalinski performed the experiments. All authors contributed to the study conception and design, material preparation and data analysis. The first draft of the manuscript was written by James F. Collawn, Rafal Bartoszewski and Magda Gebert. All authors read and approved the final manuscript.

## Acknowledgements

We would like to thank Dr. David Crossman (University of Alabama at Birmingham) for the help with GEO submission.

## Notes

### Competing Interest Statement

The authors have declared no competing interest.

