## Supplementary material for "IRE1-mediated degradation of *pre-miR-301a* promotes apoptosis through upregulation of *GADD45A*": Suplemental Figures and Table

Supplemental Figure 1

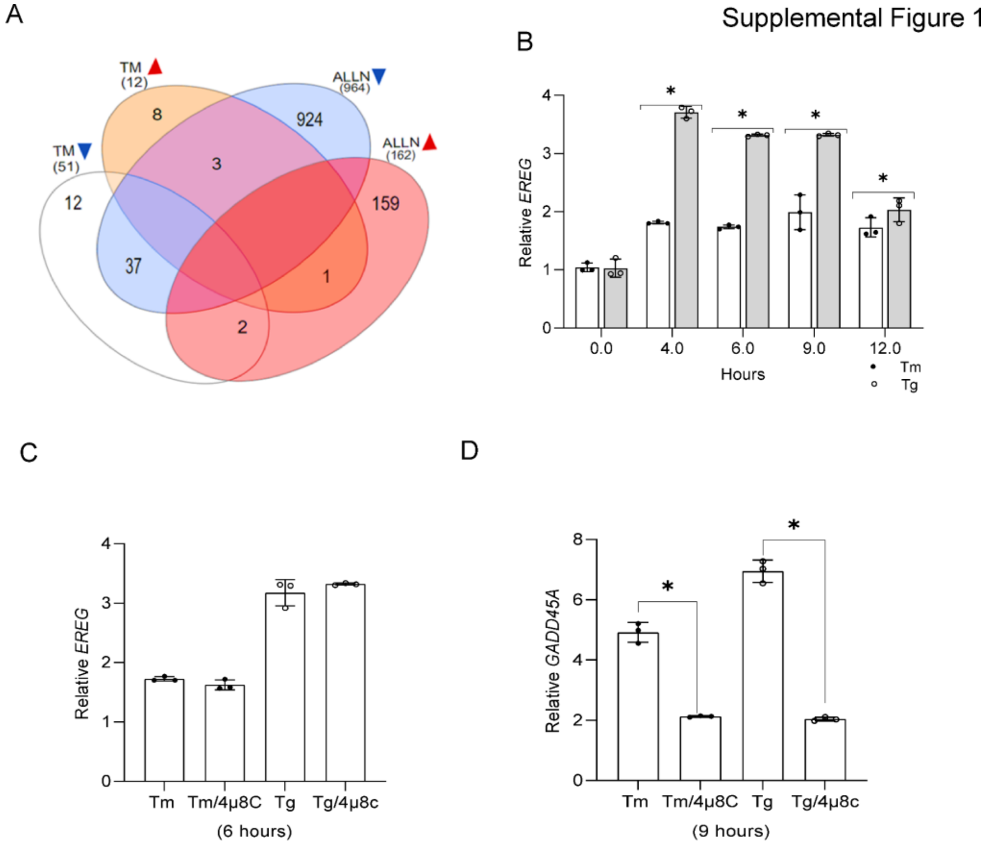


**Supplemental Figure 1. The Venn diagram** (62) **represents the general distribution of miRNAs that were significantly affected by IRE1 inhibition during ALLN and Tm induced ER stress (A).** MiRNA reduced in Tm treated 16HBE14o- cells in the presence of 4μ8C are in white the eclipse, whereas upregulated miRNAs are in light orange. MiRNA reduced in ALLN treated 16HBE14o- cells in the presence of 4μ8C are in blue eclipse, whereas upregulated miRNAs are in red. 4μ8C at 20μM concentration, Tm at 2.5μg/ml, and ALLN at 100 μM were incubated for 6 hours. ER stress-induced changes in *EREG* mRNA levels in 16HBE14o- cells for 12 hours (**B**), and the impact of 4μ8C on *EREG* expression (**C** and **D**) were analyzed by qRT-PCR and normalized to *RPLP0* mRNA levels, and expressed as a fold change over to the no-stress samples at 6 and 9 hours. The results from three independent experiments ($n = 9$) are plotted and expressed as a fold change over the no-stress controls. Error bars represent standard deviations. Significant changes ($P$ value $P < 0.05$) are marked with an asterisk. ER stressors used: Tm (2.5 μg/ml), Tg (50 nM)).

(62) Heberle, H., Meirelles, G. V., da Silva, F. R., Telles, G. P., and Minghim, R. (2015) InteractiVenn: a web-based tool for the analysis of sets through Venn diagrams. *Bmc Bioinformatics* **16**

Supplemental Figure 2

A

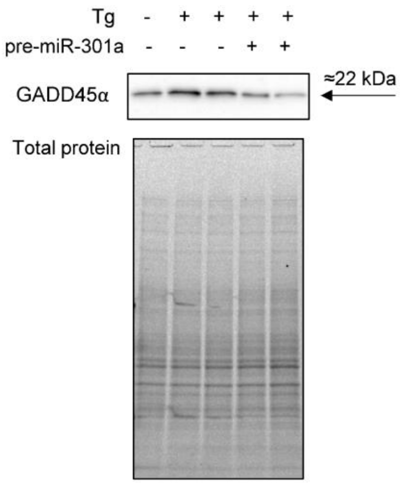


B

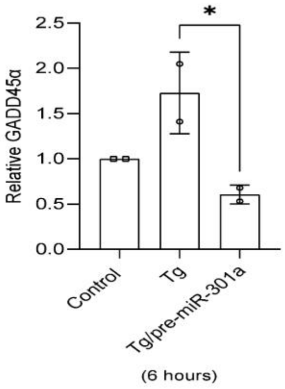


C

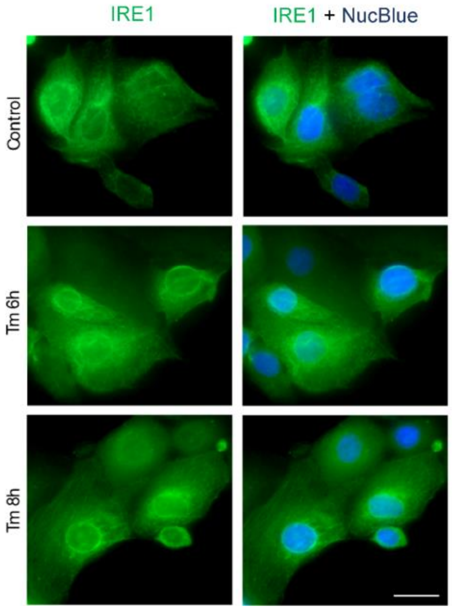


**Supplemental Figure 2. *Hsa-miR-301a-3p* affects GADD45α levels during ER stress.** 16HBE14o-cells were transfected *with pre-miR-301a* or scramble control and after 48 hours treated with Tg (50nM) for 6 h. Corresponding changes in GADD45α protein levels were monitored with Western blot (**A**) and normalized to total protein levels to the control (**B** IRE1 localizes on ER membrane-nuclear envelope interface both under both normal and ER stress conditions (2.5 µg/ml of Tm). To analyze the cellular localization of IRE1, cells were fixed with 4% paraformaldehyde and permeabilized with 0.1% Triton in PBS. Cells were blocked with 2% BSA in PBS and stained with rabbit anti-IRE primary antibodies (Abcam; #ab37073) and AF488-conjugated anti-rabbit secondary antibodies (Jason ImmunoResearch; #711-545-152). Cell nuclei were stained with NucBlue Live dye (ThermoFisher Scientific). Wide-field fluorescence microscopy was carried out using a Zeiss Axio Observer Z1 fluorescence microscope (Zeiss, Oberkochen, Germany). Images were captured using an LD-Plan-Neofluor 40 × /0.6 Korr M27 objective and an Axiocam 503 camera. A F488 signal was visualized with a 450/490 nm bandpass excitation filter and a 500/550 nm bandpass emission filter. The NucBlue Live signal was visualized using a 335/383 nm bandpass excitation filter and a 420/470 nm bandpass emission filter. Images were processed with Zeiss ZEN 2.3, FIJI and Adobe Photoshop (**C**).

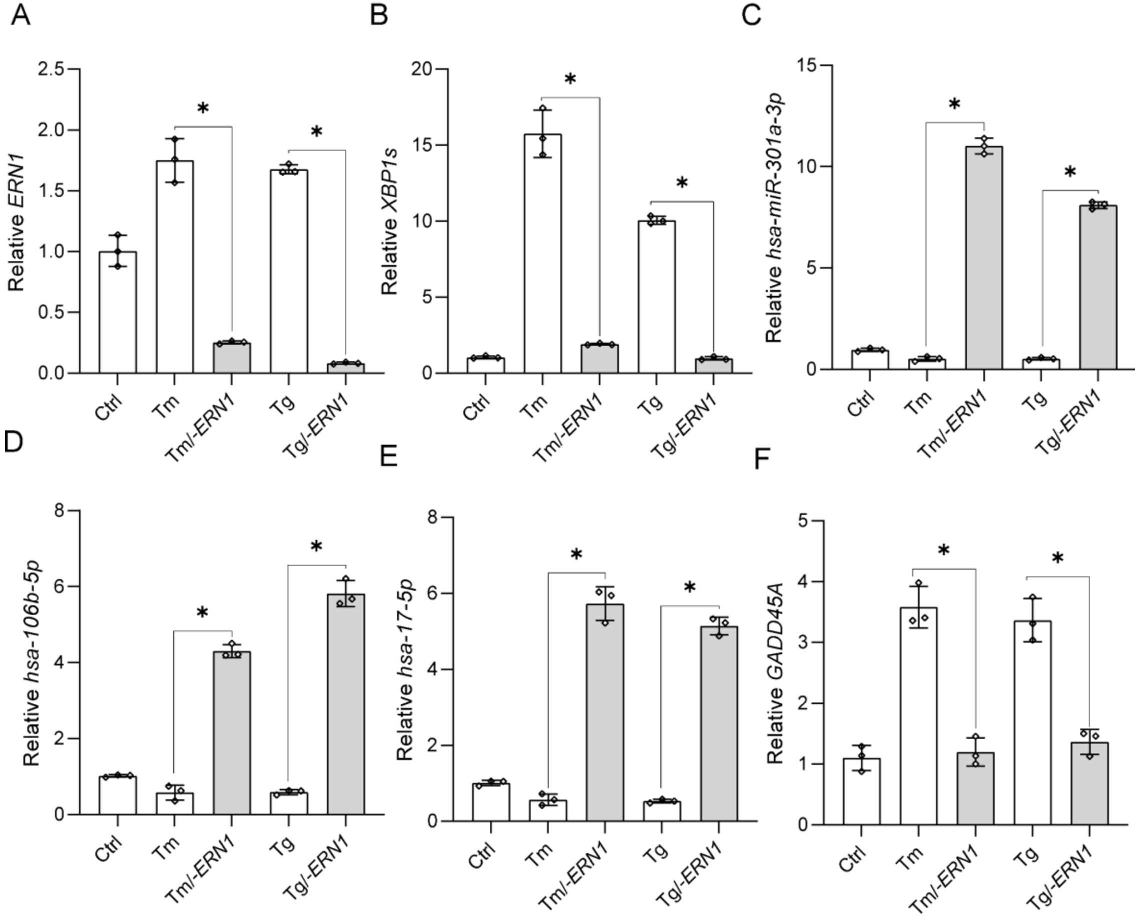


**Supplemental Figure 3. Silencing IRE1 restores *hsa-miR-301a-3p* and *hsa-miR-106-5p* expression during ER stress.** 16HBE14o- cells were transfected *with siRNA against ERN1 (*Ambion id s200432*)* or scramble control (Ambion id #4390843) (51) and after 48 hours treated with Tm (2.5µg/ml) alone or with 20 µM 4µ8C for 6 h and *ERN1*. (**A**) *XBP1s* (**B**), *hsa-miR-301a-3p* (**C**), hsa-miR-106b-5p (**D**), hsa-miR-17-5p (**E**), and *GADD45A* (**F**) RNA levels were monitored with qRT-PCR and normalized to *RPLP0* mRNA levels or *RNU44*, and expressed as the fold change over control (no induction) samples. Data represent the mean ± SD of three independent experiments (3 replicates each). * $P < 0.05$ was considered significant.

**Supplemental Table 1**. One phase decay best-fit values

| RNA | Half-life (h) | K | Tau | $R^2$ |
|---|---|---|---|---|
| *GADD45A (ctrl)* | 0.9329 | 0.743 | 1.346 | 0.859 |
| *GADD45A (Tm)* | 1.911 | 0.3627 | 2.757 | 0.812 |
| *GADD45A (Tm/4μ8C)* | 1.186 | 0.5843 | 1.711 | 0.875 |
| *GADD45A (Tm/pre-miR-301a)* | 1.414 | 0.49 | 2.039 | 0.963 |
| *pre-miR-301a (Cytosol / ctrl)* | 3.55 | 0.195 | 5.122 | 0.783 |
| *pre-miR-301a (Cytosol / Tm)* | 2.433 | 0.285 | 3.511 | 0.874 |
| *pre-miR-301a (Cytosol / Tm/4μ8C)* | 3.45 | 0.2 | 4.977 | 0.862 |
| *pre-miR-301a (Nucelus/ ctrl)* | 3.251 | 0.23 | 4.69 | 0.789 |
| *pre-miR-301a (Nucleus / Tm)* | 2.977 | 0.238 | 4.295 | 0.842 |
| *pre-miR-301a (Cytosol / Tm/4μ8C)* | 3.189 | 0.2173 | 4.601 | 0.884 |
| *pre-miR-106b (Cytosol / ctrl)* | 3.048 | 0.227 | 4.397 | 0.767 |
| *pre-miR-106b (Cytosol / Tm)* | 1.942 | 0.357 | 2.802 | 0.787 |
| *pre-miR-106b (Cytosol / Tm/4μ8C)* | 3.091 | 0.224 | 4.459 | 0.811 |
| *pre-miR-106b (Nucelus/ ctrl)* | 3.102 | 0.223 | 4.476 | 0.792 |
| *pre-miR-106b (Nucleus / Tm)* | 2.74 | 0.253 | 3.945 | 0.768 |
| *pre-miR-106b (Cytosol / Tm/4μ8C)* | 3.135 | 0.221 | 4.552 | 0.81 |
| *hsa-miR-301a-3p (ctrl)* | 9.557 | 0.072 | 13.79 | 0.774 |
| *hsa-miR-301a-3p (Tm)* | 8.513 | 0.081 | 12.28 | 0.751 |
| *hsa-miR-301a-3p (Tm/4μ8C)* | 9.488 | 0.073 | 13.69 | 0.735 |
| *hsa-miR-106b-5p (ctrl)* | 10.1 | 0.068 | 14.57 | 0.812 |
| *hsa-miR-106b-5p (Tm)* | 9 | 0.077 | 12.99 | 0.796 |
| *hsa-miR-106b-5p (Tm/4μ8C)* | 9.727 | 0.071 | 14.03 | 0.745 |
